# Sex differences in progressive multiple sclerosis brain gene expression in oligodendrocytes and OPCs

**DOI:** 10.1101/2025.02.05.636293

**Authors:** Brenna A. LaBarre, Devin King, Athanasios Ploumakis, Alfredo Morales Pinzon, Charles R.G. Guttmann, Nikolaos Patsopoulos, Tanuja Chitnis

## Abstract

Multiple sclerosis is a neurological autoimmune disease with sex-imbalanced incidence; in the USA, the disease is more likely to effect females at a ratio of 3:1. In addition, males are more likely to have a more severe disease course at time of diagnosis. Questions about both causes and downstream effects of this disparity remain. We aim to investigate gene expression differences at a cellular level while considering sex to discover fine-scale sex disparities. These investigations could provide new avenues for treatment targeting, or treatment planning based on sex.

Public single-nuclei RNA-sequencing data from three publications of progressive MS including control brains were analysed using the Seurat R package. Differential gene and pathway expression was looked at both within a specific data set which has sub-lesion level sample dissection and across all studies to provide a broader lens. This allowed for the consideration of cell types and spatial positioning in relation to the interrogated lesion in some of the calculations.

Our analysis showed expression changes in the female MS oligodendrocytes and oligodendrocyte progenitor cells compared to healthy controls, which were not observed in the corresponding male affected cells. Differentially up-regulated genes in females include increased HLA-A in the oligodendrocytes, and increased clusterin in the oligodendrocyte progenitor cells. There are also several mitochondrial genes in both the oligodendrocytes and oligodendrocyte progenitors which are up-regulated in females, including several directly involved in electron transport and which have previously been associated with neurodegenerative diseases.

These results point to altered states in oligodendrocyte progenitors and oligodendrocytes that in combination with known physiological dissimilarities between sexes may denote different programming in males and females in response to the onset of demyelinating lesions. The potential for increased debris clearance mediated by clusterin and availability of oligodendrocyte progenitors in females may indicate an environment more primed for repair, potentially including remyelination. This could contribute to the disparity in etiology in females versus males.

## Introduction

Multiple sclerosis is a common neuroinflammatory autoimmune disease with a sex-biased etiology(1–5). The disease is characterized by the presence of lesions in the brain, and one of the main McDonald diagnostic criteria for the disease is dissemination of these brain lesions in space and time, as found on MRI(6). These lesions are caused by inflammatory damage to myelin along neuronal axons. By utilizing newly developed single nuclei sequencing technologies, researchers are now able to sample cells from plaque-containing post-mortem CNS tissues(7–9) to investigate this cellular environment.

Recently, Jäkel *et al*.(8) describe various sub-groupings of oligodendroglia with altered proportions in MS vs control brains. The authors also observed changes in the normal cell balance amongst samples from normal-appearing tissue adjacent to MS lesions, including a reduction in numbers of oligodendrocyte progenitor cells (OPCs). Work by Schirmer *et al*.(9) highlighted the activated states of many cell types in MS lesions and mapped activated signatures to the rims of chronic active lesions using spatial transcriptomics. In the study by Absinta *et al*.(7), signatures of activated microglia and astrocytes were also observed with unique enrichment in the rims of chronic lesions. One innovation of the Absinta *et al*. study was also that the sampling for the single cell sequencing for these lesions occurred in several places per tissue block, so that there are separate sequencing runs for lesion core, lesion rim, and non-lesion tissue, which allows for some localization of results.

Disease severity and progression courses in MS vary by sex, with more severe cases being enriched in male subjects. The clinical outcomes of these differences have been investigated and attributed to many causes(2–5, 10), including differences in demyelination and remyelination patterns between sexes. Here, we have combined and compared single-nuclei RNA-seq (snRNA-seq) data from previously published studies to explore sex differences in gene expression across broad cell types. This approach may serve to elucidate molecular mechanisms of these known sex-related differences.

Three publicly available data sets were downloaded and processed (see Methods)(7–9). As human CNS tissue is not a readily available biospecimen, combining data sets provides us with a larger sample set to interrogate. Our integrated dataset consists of a total of 21 MS subjects (47.6% Female) and 17 Control subjects (35.3% Female), with mean ages between mid-40s and late 50s (Table 1). MS subjects were diagnosed with some form of progressive disease and had mean disease durations of approximately 20 years. Throughout this study, data from the more recent publication by Absinta *et al.*, were compared to the combined data from Jäkel *et al.* and Schirmer *et al.* to evaluate reproducibility of results.

**Table 1.** Demographics of included samples. Ages and disease durations have standard deviations in parentheses. For information on sample selection, causes of death, and IRBs, please see relevant publication. All samples for the Absinta et al. MS cohort were indicated to be “progressive”, but not whether they were primary or secondary progressive.

|  |  | Publication |  |  |  |  |  |
| --- | --- | --- | --- | --- | --- | --- | --- |
|  |  | Absinta *et al.*<sup>7</sup> |  | Jäkel *et al.*<sup>8</sup> |  | Schirmer *et al.*<sup>9</sup> |  |
|  |  | MS | Control | MS | Control | MS | Control |
| **Sex** |  |  |  |  |  |  |  |
|  | Female | 1 | 1 | 1 | 1 | 8 | 4 |
|  | Male | 4 | 2 | 3 | 4 | 4 | 5 |
| Age |  | 50.2 (6.4) | 54.7 (5.5) | 46.8 (8.4) | 58 (17.5) | 45.2 (7.0) | 53.4 (16.6) |
| **Multiple Sclerosis Type** |  |  |  |  |  |  |  |
|  | Primary |  |  |  |  |  |  |
|  | Progressive |  | - | 1 | - | 1 | - |
|  | Secondary |  |  |  |  |  |  |
|  | Progressive |  | - | 3 | - | 11 | - |
| Disease |  |  |  |  |  |  |  |
| duration |  | 19.2 (8.6) |  | 20 (6.6) |  | 19.2 (10.9) |  |

## Materials and methods

### Publicly available data

We utilized data from publicly available snRNA-seq experiments of MS samples for which fastq files were retrieved from SRA. Cell Ranger v6(11) was run to obtain count data, and then all data sets were loaded into R(12) using Seurat v3(13). Data were collected from Jäkel *et al.*(GSE118257)(8), Schirmer *et al*. (PRJNA544731)(9), and Absinta *et al*. (GSE180759)(7).

Data collection, merging and processing led to a final analyzed data sets consisting of 52,323 cells from the Schirmer *et al*. dataset consisting of 21 individuals, 12 affected and nine controls; 24,890 cells from the Jäkel *et al*. dataset consisting of nine individuals, four affected and five controls; and 58,220 cells from the Absinta *et al*. data set consisting of eight individuals, five affected and three controls.

### Data processing

Using Seurat (13), the nuclei were filtered for those that contained information from at least 500 reads, and a minimum of 200 genes but no more than 2500 genes, and those that contained < 20% reads corresponding the mitochondrial genes. Genes which appeared in less than three nuclei were also excluded.

Once cells were filtered, SCTransform(14) was used to normalize the data throughout a given sequencing run. These runs were then merged into a single Seurat object using 2000 variable features for the RunHarmony function from the Harmony(15) R package, which was applied to “harmonize” the data and account for batch effects between sample runs, different sequencing chemistries, and different data sets. To remove “cells” which may be doublets, the R package scDblFinder(16) was used.

### Cell type identification

After combining the data, clustering of the nuclei was performed. FindNeighbors was run with 13 dimensions from the harmony dimension reduction, followed by FindClusters run with a resolution of one. A UMAP was then calculated and plotted. Using known marker genes for brain cell types, each numbered cluster was than assigned one of six major cell type labels: Oligodendrocytes (OLs), OPCs, Neurons, Astrocytes, Immune, or Endothelial/Vascular.

Once these initial labels were applied, the cells underwent multiple rounds of relabeling to build confidence in the cell labels. All cells with a given label were subset from the full data, then re-harmonized and re-clustered using the same parameters as were used on the full data. Using the SingleR(17) function in R and the cell type labels provided by the authors of Jäkel *et al*., a type was assigned to every cell. The more detailed names provided by Jäkel *et al.* were collapsed down to the relevant label for the list of six given above. These labels were then used to filter the clusters. In this first pass filtering, any cluster where 95% or more of the cells were given the same label were sent on for a second round. For those clusters were there were not 95% one label, any label for which there were five or more cells were reserved to be merged later with the matching label, and any label with less than five cells had those cells removed.

The cells for a given label were now subjected to a second round of filtering. Those cells from clusters with 95% the given label, and cells from the other six cell type first round of filtering with the given label that were reserved, were brought back together and again re-merged and re-clustered, using the same parameters as the full data set. For the second filter, a cluster now had to pass a threshold of 99% the same label to avoid breakup, and any cluster that had less than 99% only had the majority cell type retained, and the rest of the cells were removed. In total, 886 cells are removed with this filtering.

Final cell numbers of twice filtered assigned labels are as follows: OLs – 72,041 cells; OPCs – 6,794; neurons –27,674 cells; astrocytes – 15,862; endothelial/vascular – 4,338; immune – 8,724 cells. For a breakdown of cells by data set, see Supplementary Table 1.

After cell type determination, each of the six major cell types underwent subtype finding. To accomplish this, the clustering step of the Seurat processing per major cell type was re-run; this does not alter the UMAP projection locations, just the cluster assignment of individual cells. A clustering resolution of 0.5 was used which resulted in several subtypes per cell type that were generally supported by the literature. Names were assigned to subtypes based on marker gene lists curated from the literature and/or pathway analyses.

## Analyses

### Differential expression

Differential expression comparisons were made using both the MS vs Control cell axis as well as cell type and sex specific axes. Using the nebula R package(18) for each comparison (the combination of compartment and sex), only the relevant cells were considered, and model formulas consisted of an intercept and the factor of interest.

### Comparison across publications

Analyses were run in three modes: only data from the Absinta *et al*. paper, data from the combined Jäkel *et al.* and Schirmer *et al.* papers, and all three data sets combined. This was done in part because of low sample numbers and to assess reproducibility across data sets. For the sex specific analyses, differential expression was evaluated for each sex in each of these modes. The results from the Absinta *et al.* only analysis were then intersected with the results from the Jäkel *et al.* and Schirmer *et al*. combined analysis, preserving those genes which were statistically significant in both (FDR < 0.05) and shared directionality in the their log fold-change (logFC). This list was then also cross-referenced with the list of differential genes generated by combining all three data sets. With the sex-agnostic analyses a similar protocol was used, with the data being run in three modes, compared between Absinta *et al.* vs Jäkel *et al.* and Schirmer *et al.*, and then checked against all three together.

### Pathway analysis

Results from the nebula analyses were then used for pathway enrichment using the fgsea R package(19). FDR values were calculated from the output results of the nebula models and used to determine the input gene lists to the fgsea function. The Canonical Pathways (CP) database from MSigDB from the Broad Institute of MIT and Harvard(20, 21) was used as the query pathways database.

### Comparison of expression levels

For comparing expression of specific genes by region, and numbers of cells by region, RNA transcript counts and cell counts were used. In the analysis comparing HLA-A expression and CD8+ T cells, the number of HLA-A transcripts were counted per compartment, and that number was divided by the number of OLs per compartment, considering only the Absinta *et al* data. The number of cells identified as CD8+ T cells as described in the methods above were also counted, and these values were then plotted in heatmaps. To compare OL and OPC marker gene expression and CLU expression, for each of the genes the transcripts were counted per region and per sex, as were the number of OL and OPC cells; the transcript count was then divided by the corresponding cell count for MBP, PLP1, OLIG1 and NG2 (labeled in the snRNA-seq data as CSPG4). For CLU, transcripts were considered from all cells types, not a specific cell subset, so transcript counts per region were divided by total cells per region, again considering only the Absinta *et al*. data for these analyses.

### Visualizations

Results from differential analyses with nebula were visualized with the R package EnhancedVolcano(22). Heatmaps were generated using the ggplot2 R package(23).

## Results

### Sex differences in gene expression by cell type

Across cell types, we observe differences in gene expression between sexes, when examining sexes individually. In females, there are significantly more differentially expressed genes in the astrocytes and OLs, compared to males, who demonstrate more expression variance in the neurons (Table 2). In the females, many of the genes with differential expression across cell types are over-expressed in the MS sample compared to control sample (Supplemental Figure 1), whereas in the males the neuronal differential expression is dominated by under-expression of genes compared to the healthy controls (Supplemental Figure 1, Supplemental Figure 2, Figure 1). Those genes which were identified by comparing the data sets were largely recapitulated in the analysis of all three datasets together, and only results which were consistent across data sets were further investigated (Table 2).

**Figure 1.**
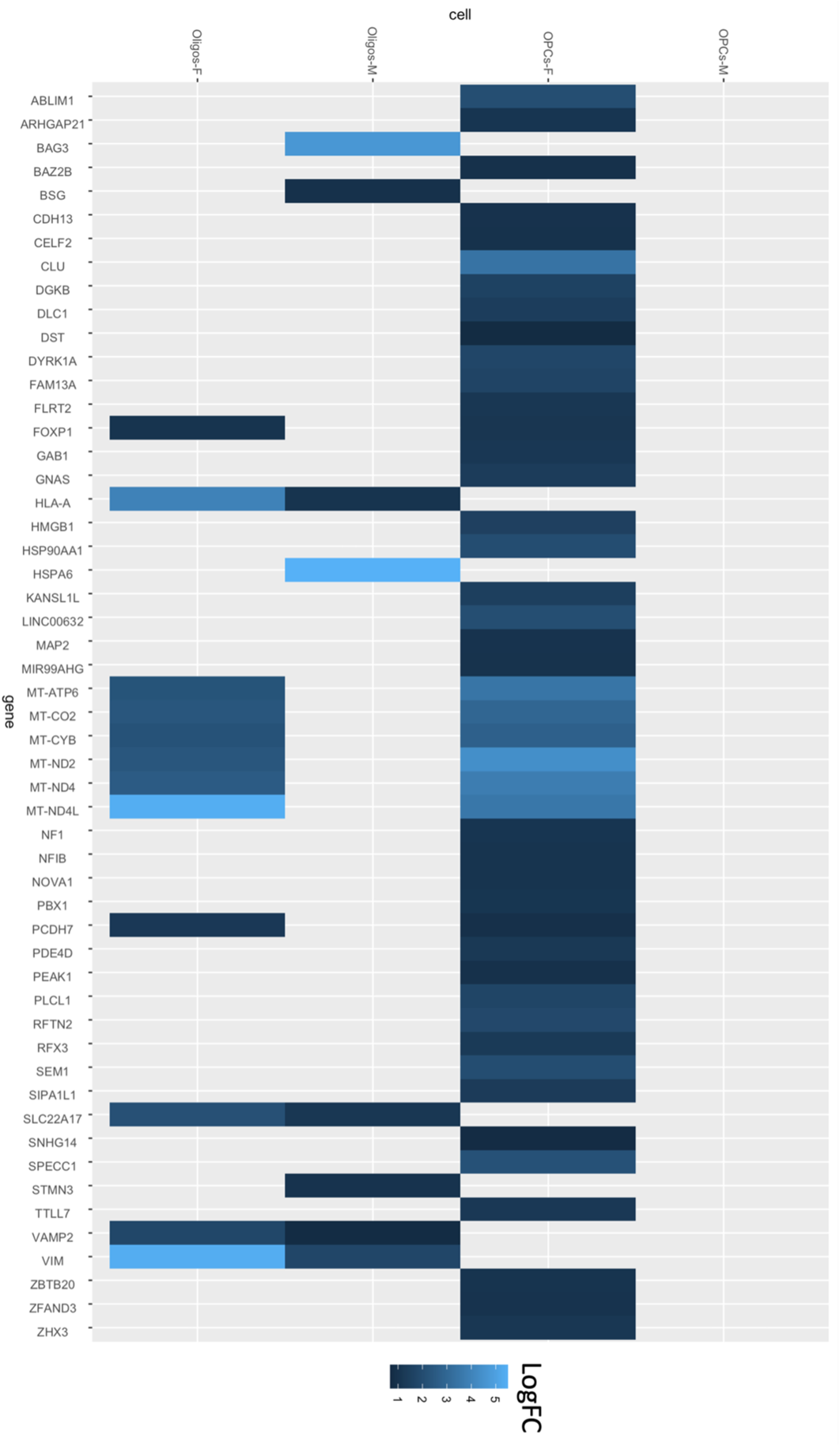
Genes which are differentially expressed in each cell subset. Genes represented are all those that were significant in male Oligos and female OPCs; this data is a subset of the data shown in the heatmap in Supplemental Figure 1; includes all genes which were significant in male OLs and female OPCs

**Table 2.** Significantly differentially expressed genes with coordinated expression between data sets. First two columns are the number of genes which were significant (FDR < 0.05) in a differential expression analysis of MS vs Ctrl cells for a given cell type in the Absinta et al. data set and were also significant in the same analysis using the combined Jäkel et al. and Schirmer et al. data set, all having log fold-changes which were the same direction. The last fourth and fifth columns are comparing the results from the first two columns to a sex-specific analysis of all three data sets, to confirm that the results from the “comparison” list weren’t artifacts Columns three and six are the non-sex-specific

| Cell | Compared between datasets |  |  | All data sets together, matching comparison |  |  |
| --- | --- | --- | --- | --- | --- | --- |
|  | Female | Male | Combined | Female | Male | Combined |
| Astrocytes | 231 | 1 | 15 | 226 | 1 | 15 |
| Endothelial | - | - | - | - | - | - |
| Immune | - | 1 | 6 | - | 1 | 6 |
| Neurons | 6 | 40 | 77 | 6 | 40 | 77 |
| Oligodendrocytes | 468 | 8 | 27 | 460 | 6 | 27 |
| OPCs | 45 | - | 1 | 45 | - | 1 |

Multiple mitochondrial genes, including MT-ATP6, MT-CO2, MT-CYB, MT-ND2, MT-ND4, and MT-ND4L were found to be upregulated in the female OPCs and OLs but not in the corresponding male cells (Figure 1). These genes are important components of the oxidative phosphorylation pathway, pointing to metabolic- or mitochondrial function-related changes in the female MS cells. While detection of mitochondrial genes can be a marker of lower-quality snRNA-seq data, or an indicator of a high proportion of apoptotic cells (24), our quality control process included steps to mitigate this (see Methods), and these changes are seen across cell types and clusters.

Another notable difference between sexes is the level of change of HLA-A in male versus female OLs. While there is significant increased expression in MS compared to control cells in both males and females, the logFC in females is 3.97 versus 1.05 in males (Figure 1). Looking at the localization of cells and expression in the Absinta *et al*. data set, we find that both the presence of CD8+ T cells and the expression of HLA-A by OLs is highest in the rim portions of the studied lesions in females, with males also having highest HLA-A expression in the rim and similar numbers of CD8+ T cells in the rim and normal-appearing brain tissue (NBT) (Figure 2). There is the least HLA-A expression and fewest CD8+ T cells present in the lesion, and an intermediate amount in the normal-appearing adjacent tissue (there were no NBT samples of female MS subjects in this data set). However, it is important to note that the numbers of cells identified as CD8+ T cells are low across samples, with only 158 cells being considered here.

**Figure 2.**
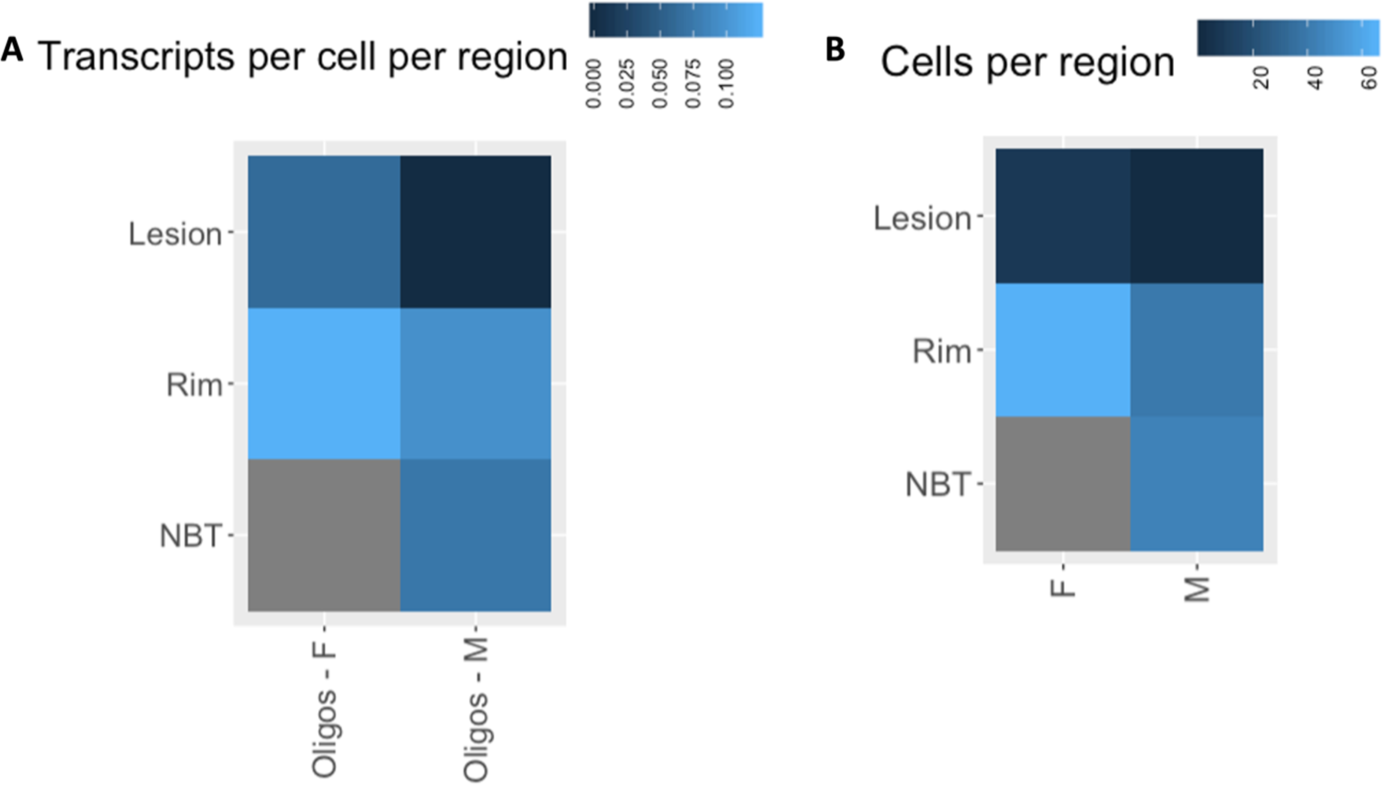
Heatmap of HLA-A transcription in OLs and CD8+ T localization by region examined in Absinta *et al*. data. For the OLs, transcripts and cells for each region were counted, and plotted values are count of transcripts divided by number of cells for each region. For CD8+ T cells, plotted values indicate number of cells identified in each region. Dark blue indicates lower values, light blue indicates higher values. NBT – normal-appearing brain tissue

### Sex-agnostic gene expression by cell type

As a second layer of evidence, and to mitigate results which may be due to the small sample sizes inherent when using only the Absinta data set, the full analysis was also done on the data without separating for sex. When performing the analyses combining all the cells of both sexes together, there is a much lower number of differentially expressed genes than in the sex-specific analysis. The exception here is the neurons, which have a greater number of significant results than in the separated analysis, perhaps due to larger sample size.

### Pathway analysis of sex-specific results

Using those genes which were coordinated across studies, a GSEA(19, 21) analysis was run using the canonical pathways (CP) database. In females, we see enrichment for different ribosomal pathways and translational pathways across results for astrocytes, OLs, and OPCs. In the neurons, we see depletion of pathways important for proper neuronal function (Supplemental Figure 3).

In OPCs, a gene contributing to many pathway results is clusterin (CLU). Clusterin was one of the genes that we find to be highly overexpressed in the MS OPCs in females, but unchanged in the male MS OPCs. This gene has previously been implicated in the lack of remyelination in Alzheimer’s disease (AD) (25), as well as apoptotic pathways and mitochondrial stabilization(26–29).

To investigate potential function of CLU in these data, the transcription of CLU across lesion compartments was compared to the expression of myelin basic protein (MBP) across compartments in both OLs and OPCs (Figure 3A, 3B, Supplemental Figure 4). MBP is a major component of the intact myelin sheaths of OL cells. Across all MS cells in Absinta *et al*. data, the CLU expression is highest in the rim, while relative MBP expression is highest in the normal-appearing brain tissue for both OLs and OPCs (Supplemental Figure 4). Looking sex-wise, MBP expression does follow a gradient for males from the NBT through the lesion core (highest -> lowest) in both OLs and OPCs, but in females the strongest MBP expression is in the rim in OLs (8.23 reads per cell in female, vs 6.91 reads per cell in male; results come from 1 female sample and 4 male). This may indicate that MBP in MS lesions is better preserved in females compared to males, either through increased resistance to demyelination or increased propensity for remyelination. Despite clear differences in the MBP expression in OLs, the OPCs show the same pattern in females as in the males. With respect to CLU, both males and females have the highest expression in the rim. Considering the differentiation of OPCs to OLs that would occur during remyelination, we also looked at three more markers, NG2, OLIG1, and PLP1(30, 31)(Figure 3C-E). OLIG1 is expressed at some level across both cell types and throughout differentiation, NG2 is expressed in OPCs and diminishes over differentiation, and PLP1, like MBP is found in mature myelinating OLs. For PLP1, we see a pattern as we would expect, more expression in the more “intact” compartments of OLs, and very low expression across OPCs. NG2 is specific to the OPCs, with the highest levels of expression in the rim cells, which is similar to OLIG1 in the OPCs; in the OLs OLIG1 expression is not as strong.

**Figure 3.**
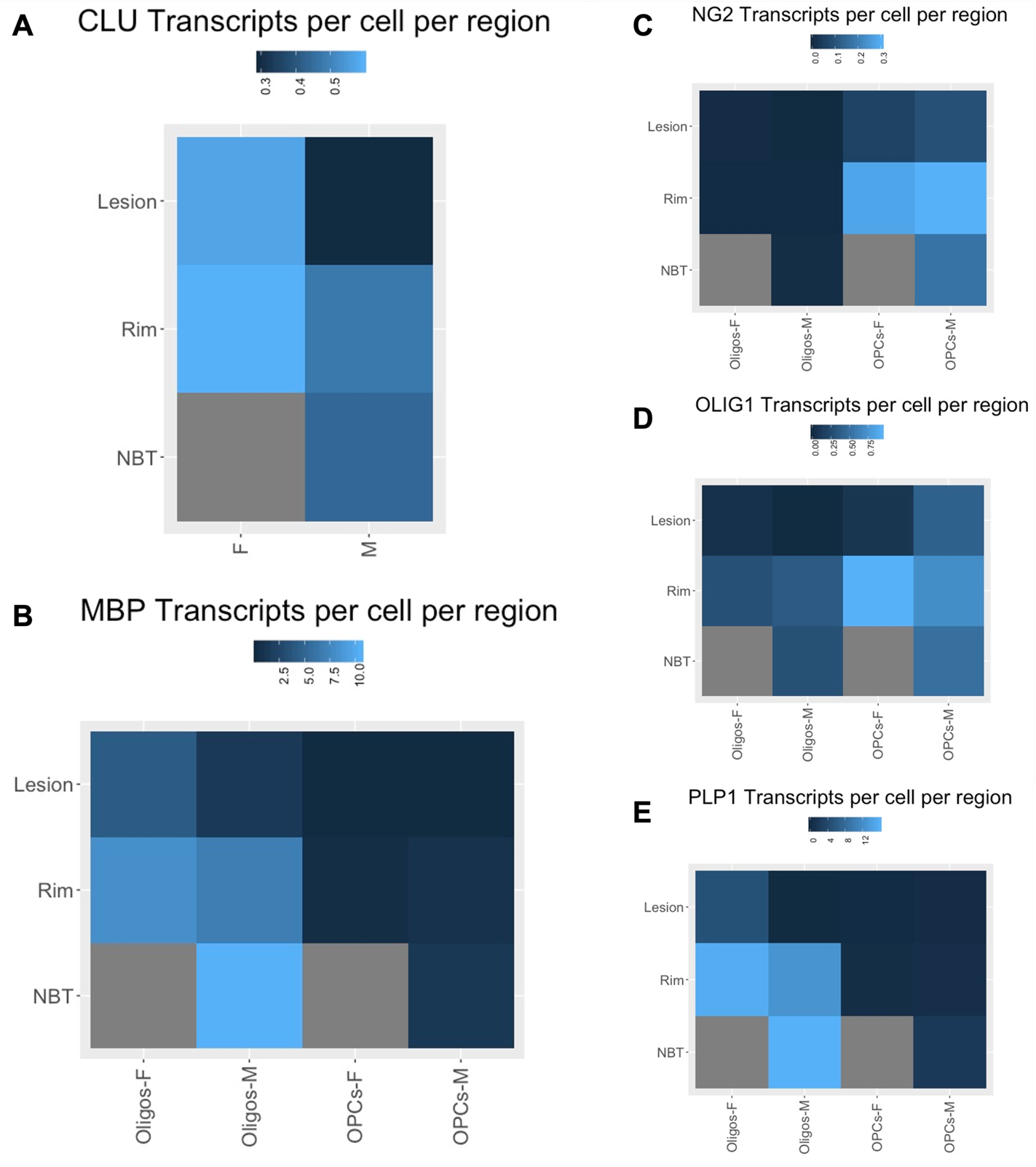
Relative expression by sex of MBP, PLP1, OLIG1 and NG2 in OLs and OPCs across lesion regions compared to CLU expression from all cells across lesion regions in Absinta *et al*. data. NBT – normal-appearing brain tissue

Among the male DEG results, we see pathways related to ribosomes across cell types, but also pathways related to different types of cellular stress (Supplemental Figure 5) including heat stress and starvation. Especially in the OPCs, there are several pathways related to heat shock factor 1 (HSF1) activation. In the OLs and astrocytes, there is increased expression of ribosome pathways, as well as translation initiation and elongation pathways, suggesting an active cellular response.

For the pathway results which are consistent between the Absinta *et al.* data set and the combined Jäkel *et al.* and Schirmer *et al.* data, there are only recapitulated results in female astrocytes, OLs and OPCs, and in male neurons. Among these results are many pathways related to other neurologic diseases (Supplemental Figure 6).

## Discussion

Our results suggest there are sex-specific differences in gene expression and pathway regulation across different cell types in MS brain lesions.

### HLA-A implicated in CD8+ T reactivity

HLA-A differential expression in OLs is an interesting result because of the genetic risk factors associated with other HLA antigens, most notably the HLA-DRB1*1501 allele, and the fact that MS is an autoimmune disease (32, 33). Though previously multiple sclerosis has been more strongly associated with MHC-class II antigens (34), HLA-A is a MHC-Class I antigen. A study in HLA-A*0201 transgenic mice which were primed with MOG peptides found a particular peptide (MOG181) that was able to stimulate a strong CD8+ T cytotoxic response; this was also shown to exacerbate MOG35-55 induced EAE (33). Similarly, using CD8+ T cells derived from multiple sclerosis affected (and control) subjects, CD8+ T cells were shown to be cytotoxic to HLA-A2 expressing OL cells even without the addition of exogenous MBP peptides (35). Thus, the increased expression of HLA-A that we observed could lead to an increase in CD8+ T cell-mediated OL loss. The higher levels of HLA-A seen in the female OLs suggest that a stronger immune response may be elicited in women.

### Clusterin in females could alter OPC differentiation, or aid in debris clearance

Clusterin has previously been investigated in the OPCs of AD patients (25). While this gene is a known risk factor for late-onset AD, it was also found to be upregulated in a subset of non-diseased mouse OPCs; upregulation in OPCs was also seen in our study. The authors found that increased phagocytosis of myelin debris and oligomeric Aβ resulted in increased CLU expression, which in turn inhibited OPC differentiation and new myelin production. Given the previously observed superior OPC activity in female versus male rodent models (4, 10), if CLU is preventing this pool of OPCs from differentiating into remyelinating OLs then this may be a driver of accumulating demyelination and downstream neuronal damage in female multiple sclerosis subjects.

However, the functional potential of CLU is complex, with many characterized and competing roles attributed to it. There exists a nuclear-located isoform promoting apoptosis(27), a mitochondrial form that averts apoptosis by preventing mitochondrial membrane permeabilization(29), and an excreted form that mainly serves as an extracellular chaperone for misfolded proteins (28). Data used here are only from mRNA, and while there are transcriptional differences between some of these isoforms which may be present in the sequencing data, those analyses were outside the scope of this study.

Considering the comparison of MBP expression and CLU expression in OLs and OPCs, the heightened expression of MBP in the female OL rim may indicate better preservation of the myelin in this region, either linked to increased resistance to demyelination or increased propensity to remyelinate. If the CLU expression in these samples indicates an apoptotic function, the rim would be where one would expect to see increased cell loss. However, studies in mice after brain ischemia showed mice which overexpressed CLU had better recovery than wild type and CLU -/- mice(36), which may indicate a high expression of CLU is beneficial to debris clearage in its role as an extracellular chaperone and may be anti-inflammatory and conducive to remyelination.

These results, in combination with the high expression of NG2 in the rim OPCs, might also indicate a stalling of remyelination at this site. As NG2 expression in OPCs goes down in the lesion compartment, this may indicate differentiation of these cells. The cells in the rim, however, maybe be pausing or accumulating before repairing the lesion damage. This could again be attributed to the CLU expression; potentially in two functions. The prevention of differentiation as seen in Beiter *et al.*(*25*) and aiding debris clearance, which may work in tandem to produce better lesion recovery over all.

These differences in OPC and OL capacity and functionality may contribute to the differences in disease course seen across sexes; females are able to recover from attacks with their greater pool of active OPCs for some time until they are disabled, whereas males have a higher initial proportion of OLs but less regenerative capacity and once they have reached critical OL loss enter a clinically recognizable and primarily progressive disease phase (3).

### Mitochondrial components are upregulated in female OPCs and OLs

Several of the pathways which were found to be recurrently dysregulated across cell types, sexes, and studies implicate oxidative phosphorylation and other mitochondrial functions. There is growing literature about the implications of mitochondrial dysfunction in multiple sclerosis. In progressive EAE models it was found that axonal mitochondrial homeostasis was disrupted prior to the symptom onset stage(37). It has also been shown that changes in mitochondrial function correlated with lesion disease course and neurological functions (38). Also, mitochondrial collapse due to the loss of the mitochondrial transmembrane potential can trigger cell death(39).

However, while altered mitochondrial function could lead to cell loss, there is also evidence that increased expression of mitochondrial genes can be triggered by the differentiation of OPCs(40), which in our data would indicate that the female OPCs are better able to differentiate than the male OPCs, and therefore lead to better recovery.

### Limitations

For this study, we used data sets produced by other laboratories that were made publicly available. While this data is a useful resource, it is a very small sample set, and in this case most of the major findings were initially identified in only one female multiple sclerosis sample (though they were corroborated with a larger data set with more female multiple sclerosis samples). Also, the snRNA-seq quality for these experiments, while sufficient for the method at the time of experiments, could now be improved upon. There is some possibility for cross-cell contamination which may affect the ability to assign a cell type to each individual sequenced “cell”. While this should be accounted for in many of the data processing methods that we employed in quality control steps, it is unlikely for all confounding noise to be eliminated.

Also, given that the nature of this study is a reanalysis of multiple data sets in a difficult to obtain, post-mortem human tissue, in vivo and/or in vitro validation of the results presented here are outside of the scope of this project; these results do however offer strong candidates for further follow-up experiments.

### Conclusions

Though this was an exploratory study based on public data, we were able to investigate sex differences in cell-specific subsets of MS brain tissue. Our results indicate that there are alterations in gene expression patterns in MS, which vary by sex and cell type. Especially in the OPCs and OLs, we see a much greater change in expression patterns in the female.

These results point to a potentially more regenerative pattern in the female compared to the male brain, leading to more aggressive clinical progression phenotypes in males and a delayed decrease in regenerative ability in females over the course of disease.

## Declarations

### Ethics approval and consent to participate

Samples and data used in this study were collected as approved in the referenced studies. IRB approval was not required for the analysis presented here.

### Consent for Publication

Not applicable

### Data availability

Data were collected from Jäkel *et al.*(<u>GSE118257</u>)(8), Schirmer *et al*. (<u>PRJNA544731</u>)(9),

### and Absinta *et al*. (<u>GSE180759</u>)(7) through the resources provided by the National Center for

Biotechnology Information (NCBI) through the National Library of Medicine at the National Institutes of Health, US.

## Funding

This work was funded by the National Institutes of Health grant R21NS123826 (to TC).

### Competing interests

Authors declare they have no competing interests. NP is now an employee of Novartis.

### Author Contributions

Authors contributed the following: BAL, NP, CRGG, TC designed the study. AP, AMP, DK collected data. BAL, DK, processed and analysed data. BAL and TC produced manuscript and figures. All authors reviewed, edited and approved the final manuscript.

## Acknowledgements

Thank you to Prashanth Sama for technical advice.

## Notes

### Competing Interest Statement

The authors have declared no competing interest.

