## Supplemental Tables and Figures for "Sex differences in progressive multiple sclerosis brain gene expression in oligodendrocytes and OPCs"

**Supplemental Table 1. Total number of cells per cell type per data set**

| | Absinta *et al.*[7] | Jäkel *et al.*[8] | Schirmer *et al.*[9] |
|---|---|---|---|
| Astrocyte | 6,843 | 1,603 | 7,416 |
| Endothelial | 1,794 | 1,776 | 768 |
| Immune | 4,029 | 1,467 | 3,228 |
| Neurons | 9,152 | 4,323 | 14,199 |
| Oligodendrocytes | 34,853 | 14,838 | 22,350 |
| OPCs | 1,549 | 883 | 4,362 |

**Supplemental Table 2. Count of all statistically significant differentially expressed genes by sex-specific comparison or all sex comparison (MS vs Ctrl), FDR < 0.05**

| | Absinta *et al.*[7] | | | Jäkel *et al.*[8] and Schirmer *et al.*[9] together | | | All three studies combined | | |
|---|---|---|---|---|---|---|---|---|---|
| **Cell** | **Female** | **Male** | **All sexes** | **Female** | **Male** | **All sexes** | **Female** | **Male** | **All sexes** |
| Astrocytes | 948 | 43 | 32 | 1062 | 443 | 372 | 2488 | 312 | 686 |
| Endothelial | 20 | 9 | 2 | 3 | 90 | 19 | 4 | 37 | 24 |
| Immune | 0 | 109 | 128 | 0 | 117 | 77 | 0 | 348 | 203 |
| Neurons | 473 | 96 | 146 | 126 | 7646 | 5459 | 278 | 7284 | 6361 |
| Oligodendrocytes | 11506 | 200 | 1242 | 530 | 1009 | 123 | 2336 | 38 | 451 |
| OPCs | 205 | 24 | 43 | 1260 | 104 | 21 | 2381 | 30 | 77 |

**Supplemental Figure 1. Volcano plots of results from sex-specific per cell type differential expression analysis in the Absinta *et al*. data.** Top row are results from females and bottom row are males; columns are Astrocytes, Endothelial/Vascular, Immune, Neurons, Oligodendrocytes and OPCs from left to right. Y axis is –log10(FDR), x axis is log2FC

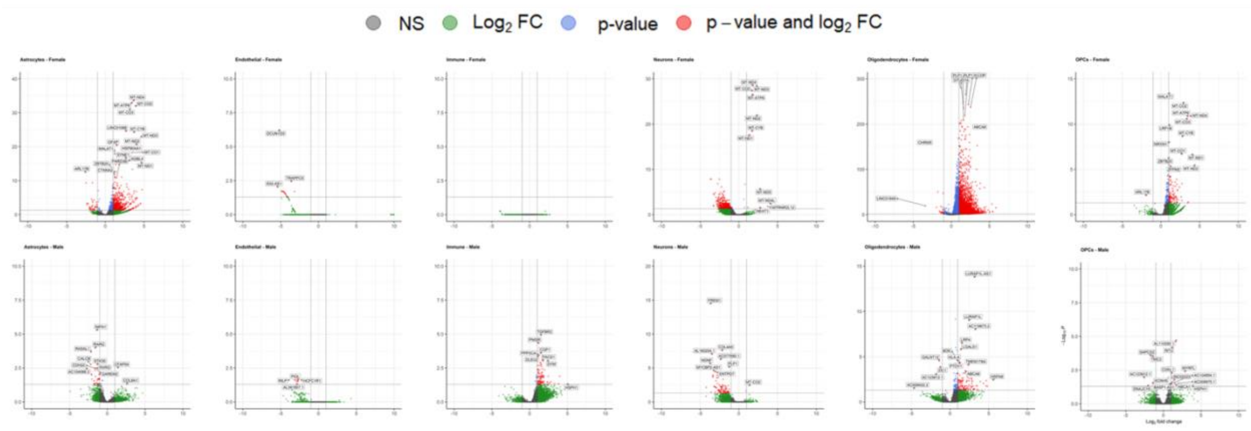

**Supplemental Figure 2. Genes which had significant differential expression (MS vs Ctrl, FDR < 0.05) per cell type per sex in Absinta *et al.* data and coordinated with results from Jäkel *et al.* and Schirmer *et al.* differential analysis per cell type per sex**. Values plotted are logFC from the Absinta *et al.* only analysis. In cell types for which there are both male and female results, we can see that many of the significant genes do not overlap.

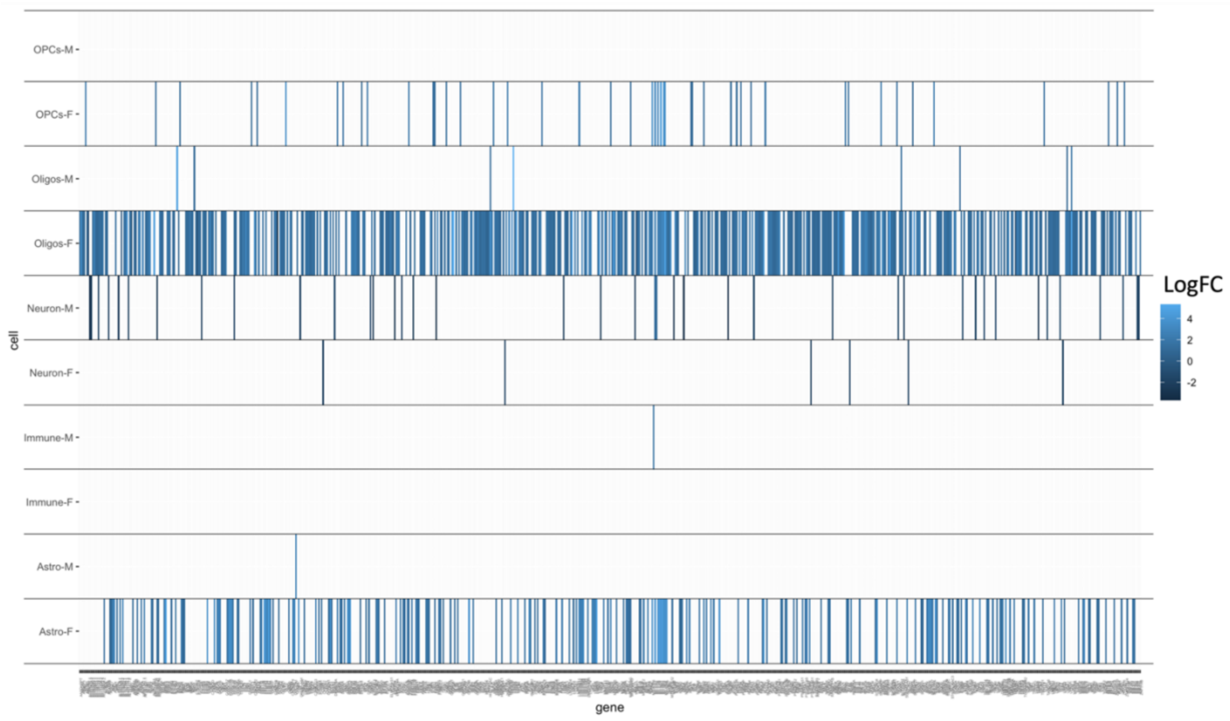

**Supplemental Figure 3. Top 10 most significant pathways (based on adjusted p-value) per cell type in female Absinta *et al.* data**. Missing cell types had no significant pathways. GSEA (gene set enrichment analysis) run using the Canonical Pathways (CP) set from the MSigDB from the Broad Institute. NES = normalized enrichment score

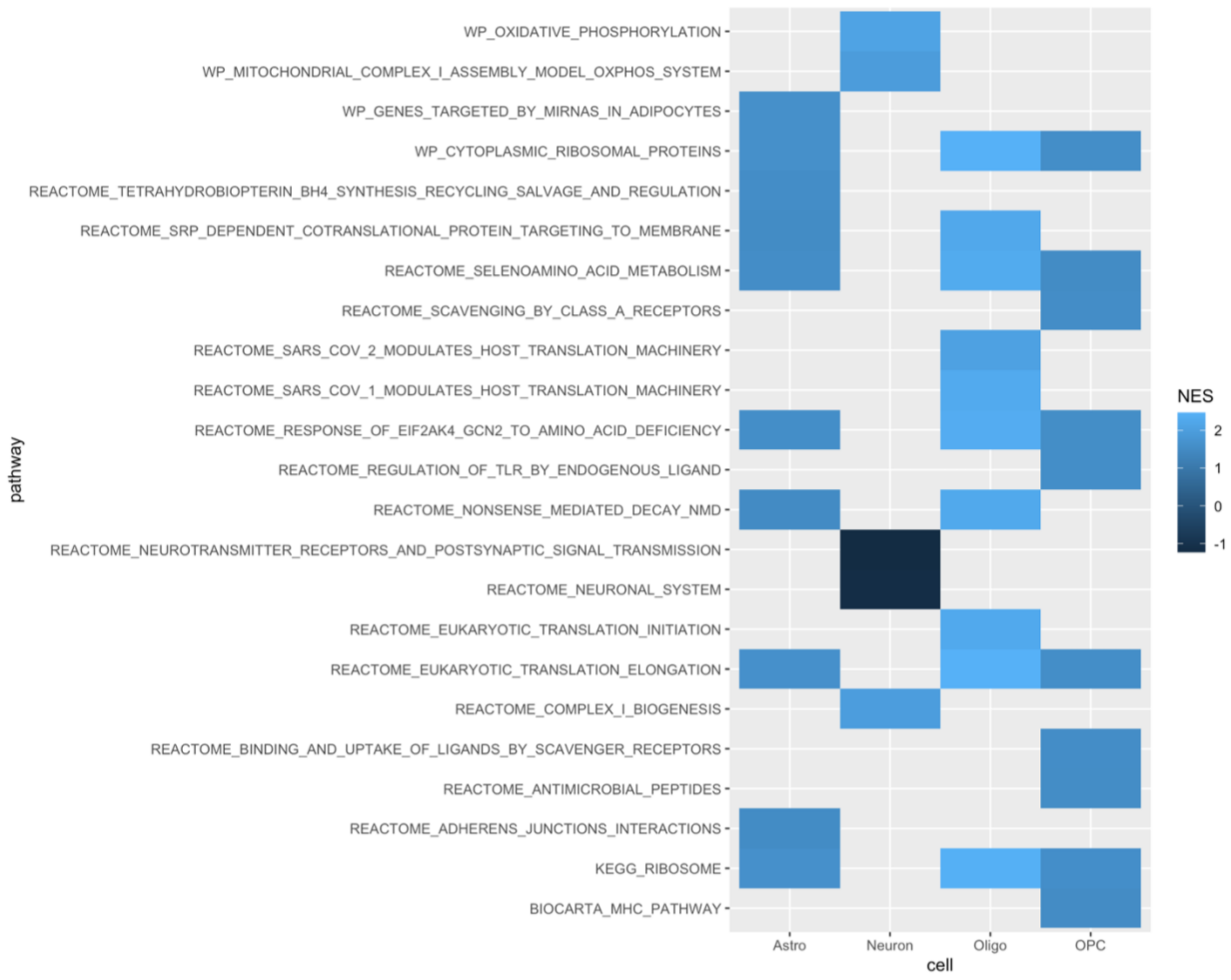

**Supplemental Figure 4. Relative expression of MBP in OLs and OPCs across lesion regions compared to CLU expression from all cells across lesion regions in Absinta et al. data.**

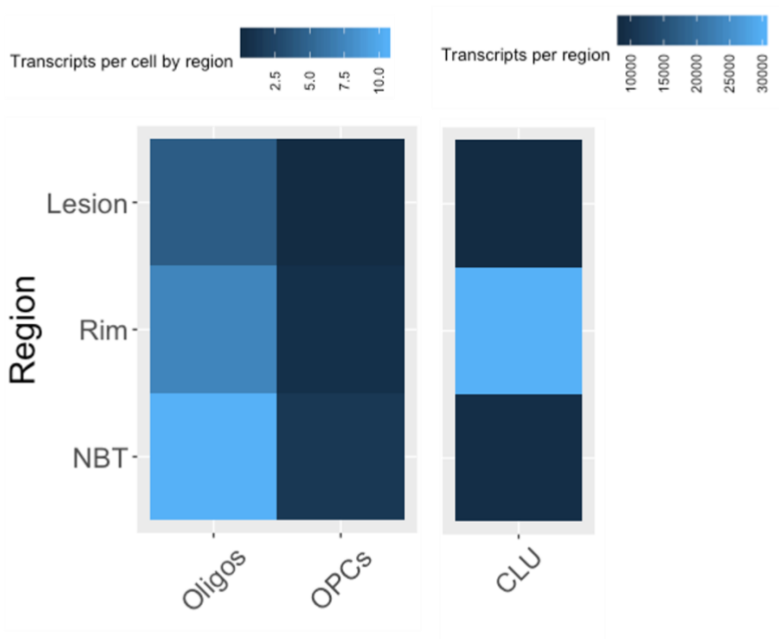

**Supplemental Figure 5. Top 10 most significant pathways per cell type (based on adjusted p-value) in male Absinta *et al.* data.** GSEA run using the Canonical Pathways (CP) set from the MSigDB from the Broad Institute. NES = normalized enrichment score

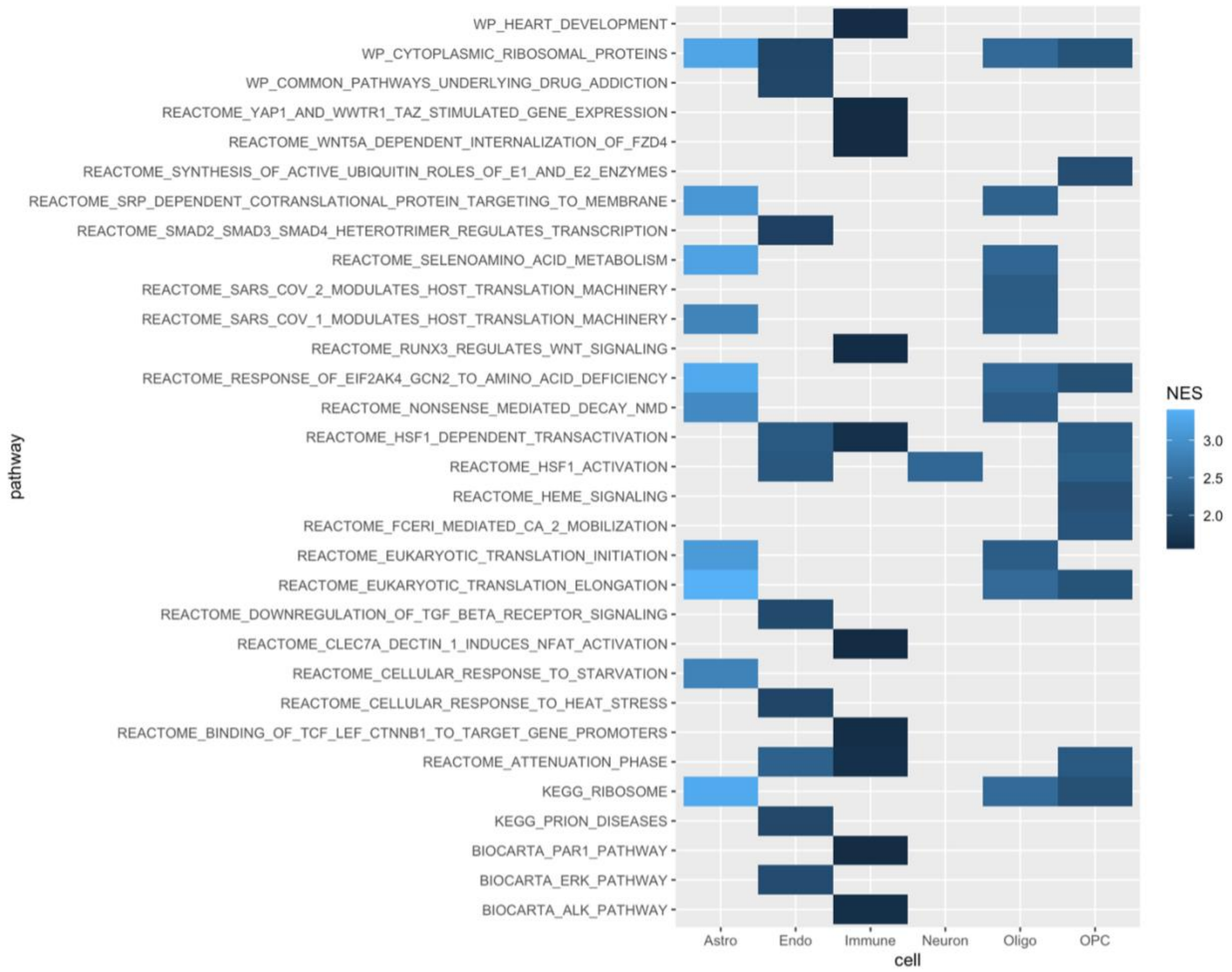

**Supplemental Figure 6. Top 10 most significant pathways per cell type (based on unadjusted p-value) in sex-specific comparisons of Absinta *et al.* vs Jäkel *et al.* and Schirmer *et al.* data.** Missing cell types had no significant pathways. GSEA run using the Canonical Pathways (CP) set from the MSigDB from the Broad Institute. NES = normalized enrichment score

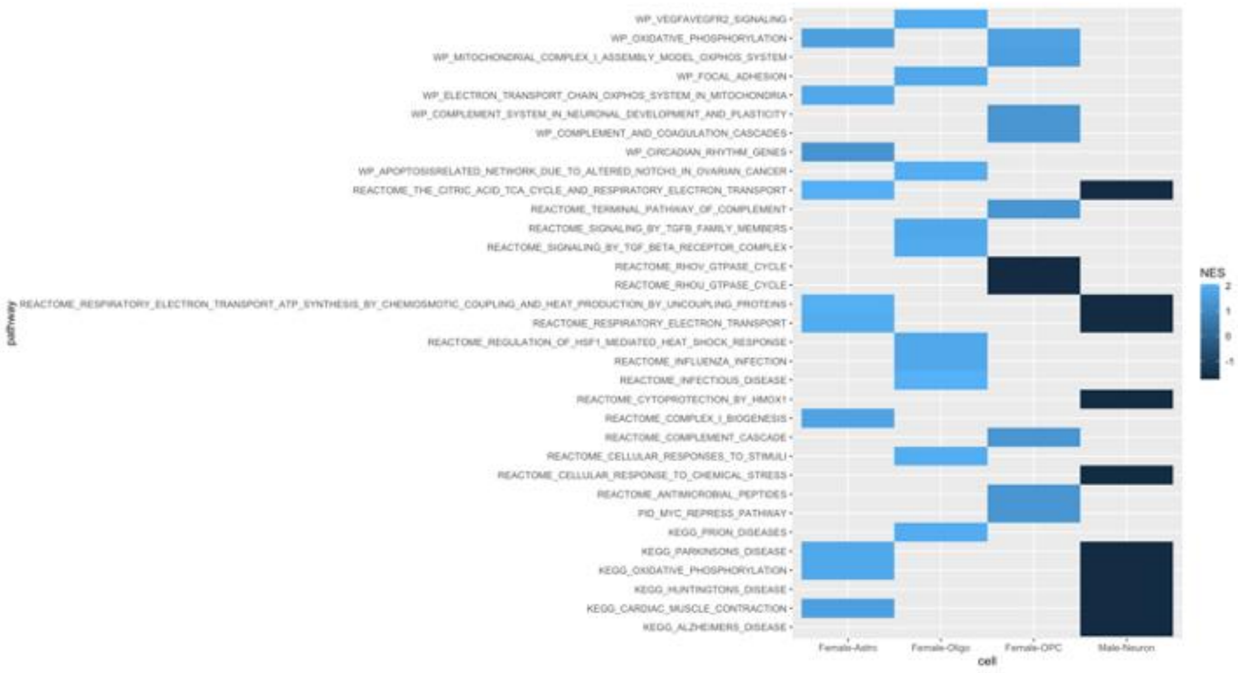
